# Common misspecification of the generation interval leads to reproduction number underestimation in phylodynamic inference

**DOI:** 10.1101/2025.04.28.649807

**Authors:** Yeongseon Park, Katia Koelle

## Abstract

Generation intervals are distributions that describe the time between infection and onward transmission. They are a key epidemiological quantity because, together with the reproduction number *R*, they determine the population-level growth rate of a pathogen and its doubling time. Conversely, when fitting epidemiological models to data, assumed generation intervals impact *R* inference. This is well-known from studies that have used case data for *R* inference, with many studies emphasizing the importance of choosing an accurate distribution for the generation interval. In phylodynamic inference of *R*, the generation interval distribution is often not explicitly mentioned, and the impact of generation interval misspecification has been studied less. Here, we explore the impact of a commonly assumed (but rarely empirically accurate) exponential generation interval distribution on the estimation of *R* in phylodynamic inference. Using simulations and inference on these simulated datasets, we find that during the early exponential growth of an epidemic, if the generation interval is assumed to be exponentially distributed when it actually has a lower variance, then estimates of *R* will be biased low. Furthermore, uncertainty in the biased *R* estimates will be small. Our work highlights the importance of acknowledging implicit generation interval assumptions in phylodynamic inference and points to the need for methodological developments in phylodynamic inference to provide greater flexibility in the specification of accurate generation intervals.

## 1 Introduction

The time between the infection of an individual and onward transmission from that individual is known as the generation interval (1). Together with the reproduction number *R*, the generation interval is key in determining how fast an infectious disease will spread through a population: a disease with a shorter generation interval will spread more rapidly through a population than a disease with a longer generation interval, provided that they have the same supercritical *R*. However, as the infection of an individual is rarely observed, this generation interval is often approximated with the serial interval (2), which is the time between the symptom onset of infector and infectee. Generation intervals also play an important role in *R* estimation. During the exponential growth of an early epidemic, reproduction numbers are generally calculated from intrinsic growth rates *r*, which could be inferred from case data to quantify the rate of change in the number of infected individuals. The calculation of *R* from *r* depends on the relationship between *r* and *R*, which is a function of the generation interval (3).

There is inherent variation in the generation interval between transmission pairs as well as between a donor and their recipients in instances where the realized number of secondary infections exceeds one. To capture this variation, the generation interval of an infectious disease is generally described with a distribution rather than by a single value. Often, the empirical shape of the distribution is not known and is difficult to estimate as infection of an individual is rarely observed (4).

In place of the empirical distribution, theoretical distributions are often used to model the generation interval. These include the gamma distribution, the Weibull distribution, and the log-normal distribution, all of which have only positive/non-negative support. This ensures that an individual does not transmit an infection prior to themselves becoming infected. The theoretical distributions that are most commonly used to model the generation interval often have high probability densities around their mean values, such that the variance of the distribution is lower than its mean. This reflects observed biological characteristics of most infectious diseases, such as secondary infections rarely occurring immediately following a primary infection. Appropriately capturing the variation in the generation interval (rather than only the mean of this distribution) is crucial when estimating the reproduction number from the epidemic growth rate, as it contributes to the quantitative relationship between *r* and *R* (3; 5).

Many models implicitly assume a generation interval distribution. In the classic susceptible-infectious-recovered (SIR) compartmental model, the generation interval is exponentially distributed, with the mean being the inverse of the rate of leaving the infectious compartment. More realistic generation interval distributions can be implemented by incorporating an additional compartment in the model that represents individuals who have been exposed (E) but have not yet become infectious. In these SEIR models, an infected individual stays in the exposed compartment before moving to the infectious compartment, delaying the time until a secondary infection can occur. This results in the generation interval distribution that is skewed toward later time-since-infection time points.

In practice, however, exponentially distributed generation intervals are often assumed, likely because of mathematical simplicity. These distributions have the highest probability density at small values and coefficient of variation of 1, generally exceeding those of empirical generation interval distributions. The assumption of an exponentially distributed generation interval (when the true generation interval distribution has a smaller coefficient of variation) is known to lead to lower estimates for *R* when using time series of case data for *R* inference (3). This is because the intrinsic growth rate r is usually estimated first from case data, and the R corresponding to this growth rate is smaller under an assumed exponential distribution than the R corresponding to this growth rate under a generation interval distribution with a smaller (< 1) coefficient of variation (3).

While misspecification of the generation interval distribution is increasingly being recognized as introducing biases in epidemiological inference based on incidence data, little attention has focused on generation interval misspecification in phylodynamic inference. In phylodynamic inference approaches, the generation interval distribution is oftentimes implicitly assumed. For example, birth-death models implicitly assume an exponentially distributed generation interval because they assume a constant birth rate (corresponding to a constant transmission rate) and a constant death rate (corresponding to an exponentially distributed infectious period).

Here, we explore the impact of generation interval misspecification on phylodynamic inference of the reproduction number *R*. Because the most common assumption in phylodynamic inference is that the generation interval distribution is exponentially distributed, we focus specifically on the impact of this misspecification when the ‘true’ (that is, empirical) generation interval distribution has a smaller coefficient of variation of less than one. Our approach relies on the simulation of mock viral sequence datasets and the subsequent application of phylodynamic inference approaches to these mock datasets to quantify the biases in *R* estimation under these approaches’ implicit assumption of an exponentially distributed generation interval.

## 2 Methods

### 2.1 Model structure for simulating mock datasets

We forward-simulated epidemiological dynamics using a susceptible-exposed-infectious-recovered (SEIR) model without susceptible depletion, which reduces to an exposed-infectious (EI) model (Figure 1A). In this model, a newly infected individual enters the exposed (E) compartment. An exposed individual either becomes infectious at rate *γ* or is sampled, which occurs at rate *ψ*. Here, we assume that the sampled individual does not transmit the infection after being sampled and thus is removed from the exposed compartment. Individuals who transition to the infectious class (I) recover from this class at rate *δ* or are removed from this compartment through sampling at rate *ψ*. In this model, an individual stays in the exposed (E) and infectious (I) compartments for an average of *L*_*E*_ and *L*_*I*_ days, respectively. *L*_*E*_ is given by 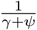 and *L*_*I*_ is given by 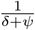. The generation interval of this model can be approximated as the convolution of two exponential distributions with means of *L*_*E*_ and *L*_*I*_ (Figure 1B). With this compartmental structure, the reproduction number *R* in the model is given by:

**Figure 1.**
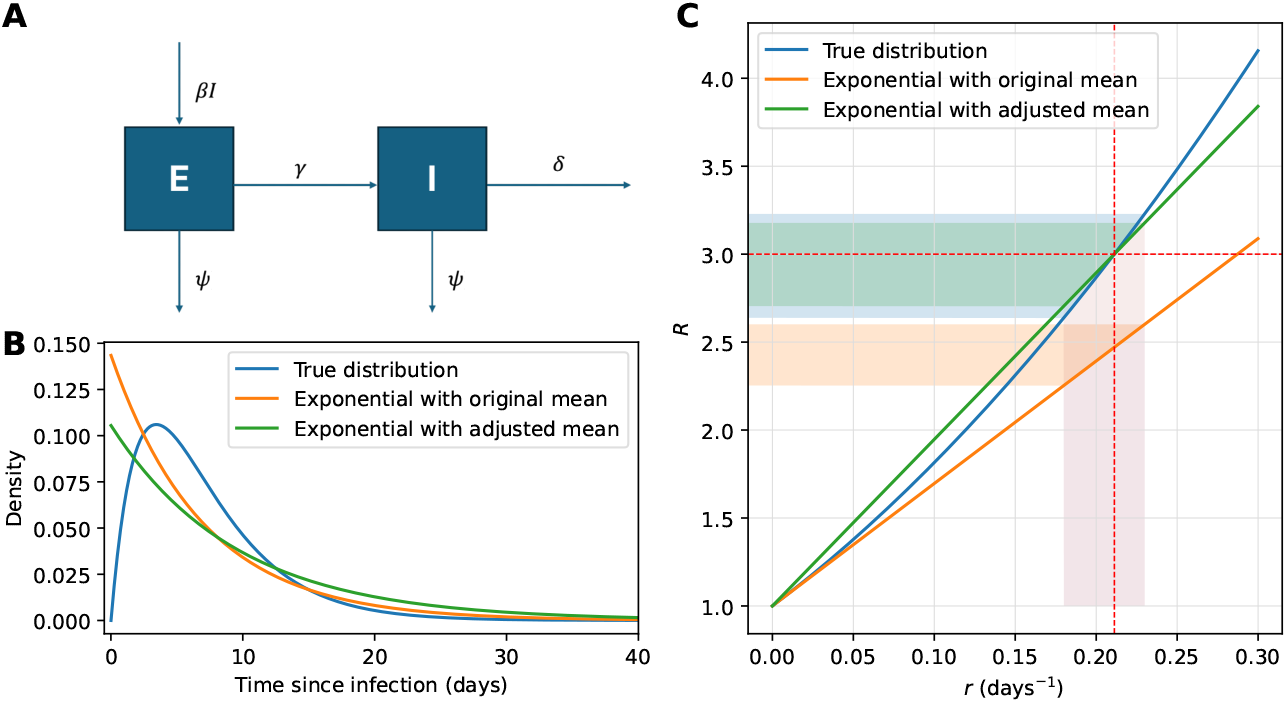
Model structure, generation intervals, and the r-R relationships. (A) Epidemiological model, consisting of exposed and infectious individuals. Individuals become infected by contact with infectious individuals and enter the exposed compartment. Individuals leave the exposed compartment by either transitioning to becoming infectious (at rate *γ*) or by being sampled (at rate *ψ*). Individuals become infectious by transitioning from the exposed class and become no longer infectious by recovering (at rate *δ*) or by being sampled (at rate *ψ*). (B) True and misspecified generation interval distributions. The true distribution is the generation interval distribution specified by the epidemiological model shown in panel A. Exponential distributions with the same mean (orange line) as the true distribution and adjusted mean (green line) that have the same *r* as the true distribution are also shown. (C) The relationship between the reproduction number *R* and the intrinsic growth rate *r* under various generation interval distribution assumptions. The red dashed lines indicate true *R* and *r* values.

**Figure 2.**
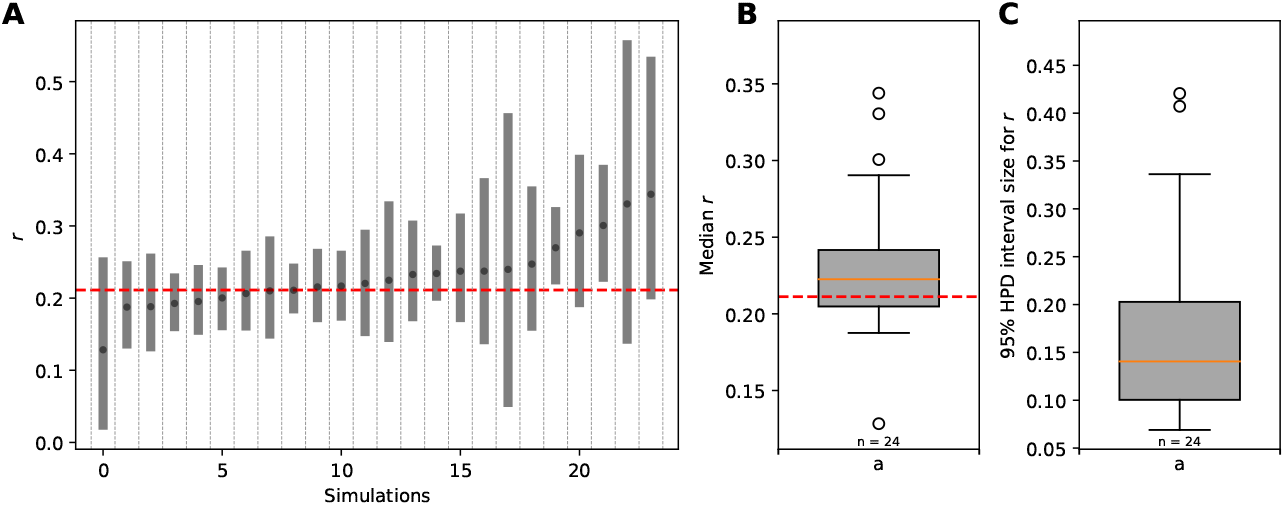
Estimated growth rates (*r*) from the exponential-growth coalescent model. (A) The median of the posterior distribution is indicated with dots, and the 95% HPD interval is shown as boxes surrounding the median. The dashed red line indicates the true *r*. Simulations are sorted by the estimated median of *r*. (B) Box plot for the median of the posterior distribution. (C) Box plot for the size of 95% HPD interval.

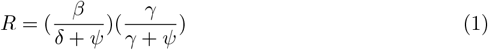

In our model, we assumed that sampling occurs from both exposed and infectious compartments so that the assumption regarding the sampling process aligns with the assumption in the constant birth-death-sampling model. The sampling probability *p*_*s*_ is calculated as:

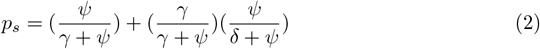

The first term captures the probability that an individual who becomes infected is sampled while in the *E* compartment. The second term captures the probability that an individual who becomes infected is sampled while in the *I* compartment. The second term includes the factor 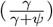 because not all individuals who become infected transition to the infectious class (some are sampled in the *E* class and therefore removed from being infected).

We simulated the epidemiological dynamics from this compartment model using Gillespie’s tau-leap algorithm (6). For the rate of becoming infectious (*γ*) and the rate of recovery (*δ*), we used 1/4 days^−1^ and 1/3 days^−1^, respectively. For the sampling rate, we used *ψ* = 0.0015 days^−1^, which, using equation (2), corresponds to the sampling probability *p*_*s*_ of approximately 1%. The per-capita transmission rate (*β*) was set to 1.01 days^−1^, which, using equation (1), corresponds to a reproduction number of 3.0. Each simulation started with an index case in the infectious (I) compartment. Simulations were run for 40 days using a *τ* of one minute.

In our stochastic epidemiological simulations, we kept track of who-infected-whom in each simulation. To simulate the evolutionary dynamics, we assigned a viral genotype to each infected individual. At the beginning of a simulation, the index case has a “reference” genotype with ancestral alleles only, and all other genotypes have a set of derived alleles relative to this reference genotype. Denoting genotypes by the set of derived alleles they carry, the reference genotype has an empty set (*G*_*ref*_ = Ø) of mutated sites. In comparison, genotype 1 (the first new genotype produced) may carry two mutated sites. These would be chronologically indexed, such that *G*_1_ = {1, 2}.

Upon infection, an individual inherits the mutated sites of the infector’s genotype plus any additional mutations that occur during the transmission event. Additional mutations would result in a new genotype. We model mutation during the transmission event as a Poisson process where the number of mutations that occur during a transmission event is drawn from a Poisson distribution with mean *p*_*m*_. We set the per-genome, per-transmission mutation rate *p*_*m*_ to 0.33 based on estimates from SARS-CoV-2 transmission pairs (7). Since we consider viral evolutionary dynamics early on during an epidemic, we assume that mutations always occur at a new site rather than hitting the same site multiple times. We, therefore, adopt an infinite sites assumption. Finally, we assume that all mutations are fitness-neutral, consistent with assumptions made in the majority of phylodynamic inference approaches.

Once a simulation has finished, viral genotypes of sampled individuals are converted to nucleotide sequences for downstream BEAST analyses. Ancestral alleles for each site are first randomly chosen from among the four nucleotide bases (‘A’, ‘T’, ‘G’, and ‘C’). The reference genotype is set as the genome carrying all ancestral alleles. Derived alleles are then randomly chosen from among the remaining three nucleotide bases at each site. If the viral genotype of an individual *G*_*i*_ has site *j* as an element, virus *i* then has the derived allele at site *j*. (If site *j* is not included in the viral genotype, then virus *i* has the ancestral allele at the site.)

### 2.2 Bayesian phylodynamic analyses

All phylodynamic analyses were performed using BEAST v. 2.7.5 (8) using three different models: (1) the exponential growth coalescent model (9), (2) the multi-type birth-death model implemented in the BDMM Prime package (10), (11), and (3) the PhyDyn coalescent model (12). The BDMM Prime is the extended version of the original BDMM package (13), which is for multi-type birth-death models with migration (13). This model can incorporate structured populations by considering each type as a “subpopulation” or a “deme” and transition between each type as “migration” (13). The PhyDyn model is a coalescent-based model that allows for complex population dynamics as well as population structure using generalized coalescent rates (14).

For each of these three models, we specified the same model of sequence evolution: the Jukes-Cantor substitution model (15) and a strict molecular clock. For each model, we calculated the reproduction number from the estimated parameters of the model. For example, for the exponential growth coalescent model, we calculated *R* from the exponential growth rate, and for the birth-death model and for PhyDyn, we calculated *R* from the estimated transmission rate. We obtained the median and 95% HPD of the calculated reproduction numbers using a custom script (available on https://github.com/yspark576/misspecified_generation_interval). All BEAST MCMC chains were run sufficiently long to result in effective sample size (ESS) values larger than 200 for the parameters of interest. ESS values were calculated using LogAnalyser within the BEAST2 package. Convergence of the MCMC chains was further assessed using Tracer v. 1.7.2 (16).

To investigate the impact of generation interval misspecification on phylodynamic inference, we first performed each of these three phylodynamic analyses under different assumptions of the generation interval. First, to assess the bias introduced by a misspecified exponential distribution, we compared the estimation under (1) the true generation interval distribution and (2) the generation interval distribution that has the same mean but is exponentially distributed. We further explored the results under (3) an exponential distribution parameterized with a mean generation time that would reproduce the true intrinsic growth rate *r* of the epidemic (Figure 1C). We performed this latter analysis to determine whether a misspecified exponential distribution would yield unbiased estimates of *R* when the mean of this distribution was set to a value that would reproduce the correct epidemic growth rate.

#### 2.2.1 Phylodynamic analysis under the coalescent exponential model

The coalescent model with exponential growth does not directly estimate the reproduction number *R*. Instead, the model estimates the intrinsic growth rate *r*. The estimate of *r* is then used to calculate the estimated *R* using the *r*-*R* relationship provided in (3). This calculation requires the specification of the generation interval distribution. Under the ‘true’ generation interval distribution of our exposed-infectious (EI) compartment model (Figure 1), the *r*-*R* relationship is given by (3):

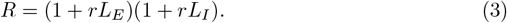

Given our *δ, γ*, and *ψ*, the *L*_*E*_ and *L*_*I*_ were 3.976 and 2.987 days, respectively, close to the values of 4.0 and 3.0 days, respectively, which would have been the case in the absence of sampling. We, therefore, use this relationship to calculate *R* from estimated *r* values under generation interval assumption (1) (the true distribution). Under an exponential generation time distribution, the relationship between *r* and *R* is given by (3):

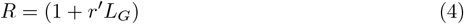

where *r*^*′*^ is the growth rate under an exponentially distributed generation interval and *L*_*G*_ is the mean generation interval, which is given by *L*_*E*_ + *L*_*I*_ = 6.963 days. We use this relationship to calculate *R* from estimated *r*^*′*^ values under generation interval assumption (2) (the exponential distribution parameterized with the true mean generation interval *L*_*G*_). For generation interval assumption (3), we used the same equation (4) but with the mean generation interval adjusted to match the true intrinsic growth rate. This adjusted mean 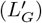 is calculated by setting *r* = *r*^*′*^, and solving for *L*_*G*_. Under *R* = 3, the adjusted mean generation interval is calculated to be 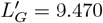 days.

#### 2.2.2 Phylodynamic analysis under the birth-death model

The true generation interval from the EI model structure (assumption 1) was modeled using the multi-type birth-death model implemented in the BDMM-Prime package for BEAST2. In this multi-type birth-death model, the exposed (*E*) and infectious (*I*) compartments were considered as separate “demes”. Transmission is considered as a “birth” from deme *I* to deme *E*. Individuals who become infectious “migrate” from deme *E* to deme *I*, and recovery of an individual corresponds to a “death” from deme *I*. Consistent with the structure of our simulation model, individuals in demes *E* and *I* can be sampled. Individuals, once sampled, are removed from being considered infected. We set the migration rate from deme *E* to *I* to *γ*, the death rate from deme *I* to *δ*, and the sampling rate from demes *E* and *I* to *ψ*. We estimated the birth rate from deme *I* to deme *E*, which corresponds to *β*, along with the time to the index case (*T*_0_), which is the time between the last sample date and the infection of the index case. We used a uniform prior distribution for *β* with lower and upper bounds of 0.337 and 3.368 days^−1^, respectively, corresponding to lower and upper bounds on *R* of 1 and 10, respectively. For the time to the index case (*T*_0_), the lower bound for the uniform prior from 0 to infinity, where the timing of the index case could be anytime before the last sample date. For the analyses, we further provided accurate information about the deme (*E* or *I*) from which each individual was sampled.

The exponentially distributed generation interval distributions (assumptions 2 and 3) were implemented using a single-type birth-death model by setting the number of demes to 1 in the BDMM-Prime model. In the single-type birth-death model, each individual stays infectious for 1*/*(*δ* + *ψ*), and thus, the mean generation interval from the model is 1*/*(*δ* + *ψ*). Also, the sampling proportion *p*_*s*_ is fixed to the same value used in the true model. Based on this, the death rate *δ* and the sampling rate *ψ* for the model were calculated from 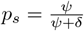 and *ψ* + *δ* = 1*/L*_*G*_, as *ψ* = *p*_*s*_*/L*_*G*_ and *δ* = (1 − *p*_*s*_)*/L*_*G*_. These parameters were then fixed at these calculated values in the analyses.

For assumption (3), 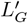 was used instead of *L*_*G*_ to calculate the death rate *δ* and the sampling rate *ψ*. As for assumption (1), we estimated the transmission rate *β* and the time to the index case (*T*_0_). We used a uniform prior for *β* with lower and upper bounds, respectively, corresponding to *R* values of 1 and 10, using the equation *β* = *R*(*δ* + *ψ*). For the time to the index case, we used the same uniform prior as for assumption (1). Table S1 summarizes the priors used in the BDMM-Prime model under all three generation interval distribution assumptions.

#### 2.2.3 Phylodynamic analysis under a coalescent model with complex dynamics

To incorporate the EI model under a structured coalescent framework, we used the PhyDyn package implemented in BEAST2 (12). As in the multi-type birth-death model, under the assumption of true generation interval distribution, transmission is modeled as a birth event from deme *I* to deme *E*, the transition from exposed to infectious compartments is modeled as a migration event from deme *E* to deme *I*, and recovery from the infectious class is modeled as a death from deme *I*. Sampling events from compartments *E* and *I* are modeled as death events.

As in the birth-death analyses, the values of parameters *ψ, γ*, and *δ* were fixed at their true values, and we estimated *β*. For *β*, uniform priors with lower and upper bounds that correspond to the *R* of 1 and 10, respectively, were used. As before, the compartment from which each sampled individual was sampled was specified. The exponential generation interval distributions were implemented with a single infected compartment where individuals become no longer infectious, either through recovery or sampling, with rates 1*/L*_*G*_ and 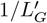, respectively, for generation interval distribution assumptions (2) and (3). With the recovery rate fixed, the per-capita transmission rate *β* was estimated. Table S2 summarizes the priors used in the PhyDyn model under all three generation interval distribution assumptions. Note that sampling is not explicitly incorporated in PhyDyn models. It is considered as a removal from the compartment.

## 3 Results

Below, we first compare the estimated *R* under the true distribution and misspecified exponential distribution to determine whether the misspecification leads to bias in phylodynamic inference (indicated with blue and orange in Figures 3, 4, and 5). In the following section, we further investigate whether the bias we observe with a misspecified exponential can be explained by the *r* − *R* relationship through the exponential distribution with adjusted mean (indicated with green in Figures 3, 4, and 5).

**Figure 3.**
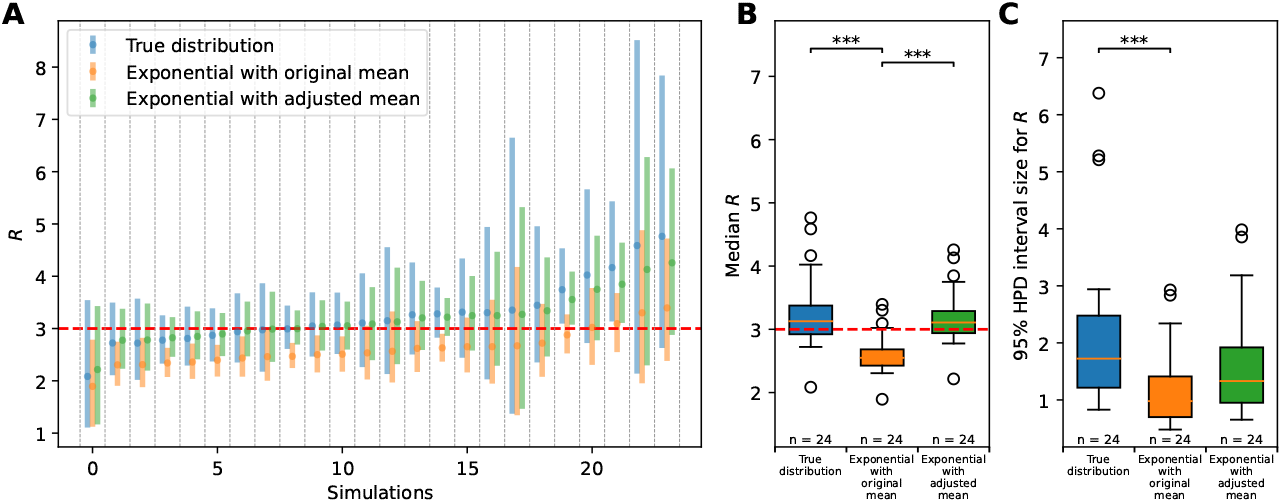
Estimates of *R* from the exponential-growth coalescent model with different generation interval distributions. (A) The median of the posterior distribution is indicated with dots, and the 95% HPD interval is shown as boxes surrounding the median. The dashed red line indicates the true *R*. Simulations are shown as ordered in Figure 2A. (B) Box plot for the median of the posterior distribution. (C) Box plot for the size of 95% HPD interval. For (B) and (C), the asterisk indicates the p-value for the Mann-Whitney U rank test (*p <* 0.001 for ‘***,’ *p <* 0.01 for ‘**,’ and *p <* 0.05 for ‘*’) comparing two distributions.

**Figure 4.**
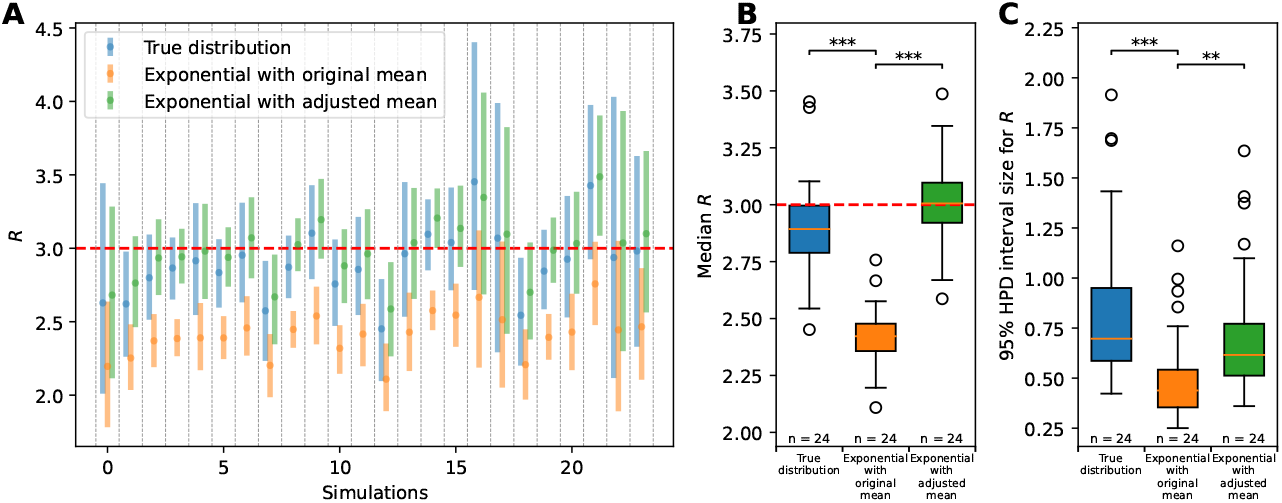
Estimates of *R* from the BDMM model with different generation interval distributions. (A) The median of the posterior distribution is indicated with dots, and the 95% HPD interval is shown as boxes surrounding the median. The dashed red line indicates the true *R*. Simulations are shown as ordered in Figure 2A. (B) Box plot for the median of the posterior distribution. (C) Box plot for the size of 95% HPD interval. For (B) and (C), the asterisk indicates the p-value for the Mann-Whitney U rank test (*p <* 0.001 for ‘***,’ *p <* 0.01 for ‘**,’ and *p <* 0.05 for ‘*’) comparing two distributions.

**Figure 5.**
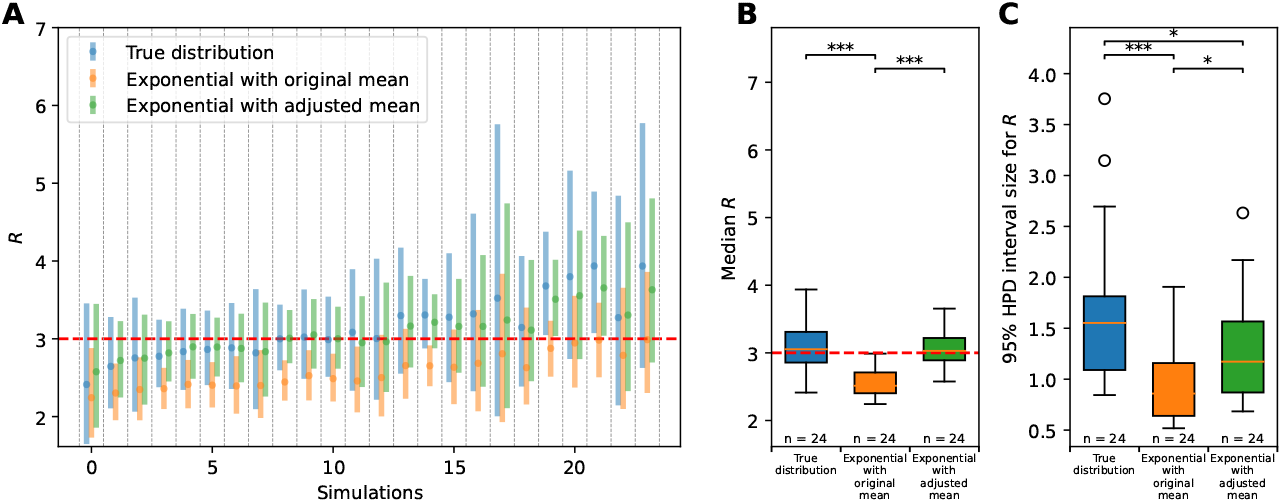
Estimates of *R* from the PhyDyn model with different generation interval distributions. (A) The median of the posterior distribution is indicated with dots, and the 95% HPD interval is shown as boxes surrounding the median. The dashed red line indicates the true *R*. Simulations are shown as ordered in Figure 2A. (B) Box plot for the median of the posterior distribution. (C) Box plot for the size of 95% HPD interval. For (B) and (C), the asterisk indicates the p-value for the Mann-Whitney U rank test (*p <* 0.001 for ‘***,’ *p <* 0.01 for ‘**,’ and *p <* 0.05 for ‘*’) comparing two distributions.

### 3.1 *R* is systematically underestimated under a misspecified exponential distribution with true mean

#### 3.1.1 Coalescent exponential growth model

We simulated 30 mock datasets with sample size ranges from 11 to 310 and attempted to first estimate the intrinsic growth rate *r* using the coalescent exponential growth model implemented in BEAST2 (Figure 2) for each of these datasets. Coalescent inference with 6 of the 30 mock datasets did not reach convergence even after 24 million chains. These six datasets each had a small sample size and contained little genetic variation. Below, we therefore focus on the 24 simulations where effective sample sizes exceeded 200. As we generated our simulated datasets using the EI model with an *R* of 3.0, the exponential growth rate is expected to be 0.211 per day from Equation 3.

Of the 24 datasets, 22 had the true value of *r* contained within their 95% HPD (Figure 2A). The median estimated *r* values were slightly higher than the true value of *r* = 0.211 per day, with a bias toward higher median growth rates. This bias was statistically significant (one-sample Wilcoxon rank test, *p <* 0.05, Figure 2B) and is consistent with the push-of-the-past effect (17). (Stochastic simulations of an emerging epidemic can result in various outcomes, including extinction. However, we only considered the replicates with surviving epidemics, which tend to have a “flying start”. As a result, the effective growth rate from those simulations is higher, which can lead to the overestimation under the deterministic assumption of the coalescent model.)

We converted these *r* estimates into *R* estimates using equation 3 for the true generation interval distribution and using equation 4 for the exponential generation interval distribution with the true mean *L*_*G*_. Under the true generation interval distribution (assumption 1), the true *R* was included in the 95% HPD interval in 22 out of 24 datasets (Figure 3A). As expected, the datasets that failed to recover the true *R* were the datasets that failed to recover the true *r*. The median of the estimated *R* was slightly higher than the true value (one-sample Wilcoxon rank test, *p* = 0.041, Figure 3 B), reflecting the slight bias in estimates of growth rates.

Under the exponential distribution with the same mean as the true distribution, however, the median of the *R* estimate was significantly lower than the true value (one-sample Wilcoxon rank test, *p <* 0.001), and the true *R* was included in the 95% HPD of only 12 of the 24 datasets. In the remaining datasets, *R* was underestimated, and the 95% HPD failed to capture the true value. This suggests that even when the true *r* was recovered, *R* tends to be underestimated under a misspecified exponential distribution parameterized with the true mean. These results are consistent with findings based on case data (3; 5). This underestimation could be explained by the higher growth rate under the exponentially distributed generation interval (Figure 1C). To match the estimated growth rate from the data, the model uses an exponential distribution assumption, which leads to a lower *R* estimate, thereby compensating for the distribution’s naturally higher growth dynamics.

In addition to the underestimation of *R* itself, the sizes of the 95% HPD interval estimates were also smaller under the misspecified exponential distribution compared to the true distribution (Mann-Whitney U rank test, *p <* 0.001, Figure 3C). This could also be explained by the *r* − *R* relationship under different generation intervals (Figure 1C). Given the lower and upper bounds of the 95% HPD interval for *r* as (*r*_*l*_, *r*_*h*_), we can approximate the width of the 95% HPD interval for *R* by calculating *f* (*r*_*h*_) − *f* (*r*_*l*_), since *R* = *f* (*r*) is a monotonically increasing function. Since the derivative of *R* = *f* (*r*) is greater for the true distribution compared to the exponential distribution, *f* (*r*_*h*_) − *f* (*r*_*l*_) is greater for the true distribution, and thus the uncertainty is underestimated under the exponential distribution. The underestimate of uncertainty in estimates with a misspecified exponential distribution emphasizes that careful interpretation is crucial under the misspecified generation interval.

#### 3.1.2 Birth-death model

Unlike in the exponential-growth coalescent model, where *r* is estimated and the generation interval distribution is only used thereafter to convert *r* to *R*, the birth-death model incorporates the generation interval distribution into the inference itself. Therefore, separate analyses were performed for each of the three generation interval distribution assumptions using the same simulated datasets. In Figure 4, we show the same data set as shown in Figure 3 for the exponential-growth coalescent model.

For the birth-death model, to obtain the estimate for *R*, we converted the sampled *β* into *R* using equation 2.1 and obtained the median and 95% HPD interval. Among the 24 datasets, 20 datasets had 95% HPD interval that successfully captured the true value (Figure 4A). The four datasets that failed to capture the true value underestimated the reproduction number. The median estimated *R* values were slightly lower than the true value (one-sample Wilcoxon rank test, *p* = 0.047) with a mean of 2.896 (Figure 4B).

Under the exponentially distributed generation interval, implemented as a single deme, the true values were captured in only 4 of the 24 datasets (Figure 4A). The median of the estimated *R* was significantly lower than the true value one-sample Wilcoxon rank test, *p <* 0.0001) with a mean of 2.412 (Figure 4B). This suggests that *R* is underestimated even when the *r* − *R* relationship was not used directly to estimate the reproduction number as in the exponential growth coalescent model. Even when the generation interval is indirectly assumed in the model, the generation interval distribution affects the estimation of the reproduction number.

As in the exponential growth coalescent model, the size of the 95% HPD interval was also smaller under the exponential distribution (Mann-Whitney U rank test, *p <* 0.001; Figure 4C), with a mean of 0.858 and 0.513 for the true and exponential distribution, respectively. This again indicates that misspecification of the generation interval distribution using an exponential distribution will bias estimates of *R* to be low and that the uncertainty in these estimates will also be too low.

#### 3.1.3 Coalescent model with complex dynamics (PhyDyn model)

Similarly to the birth-death model, the PhyDyn model incorporates the generation interval distribution into the inference itself. Separate analyses were therefore performed again for each of the three generation interval distribution assumptions using the same simulated datasets. Figure 5 shows the same 24 simulated datasets from Figures 3 and 4.

As in the BDMM model, we obtained samples of the transmission rate *β* from the MCMC chains and converted them to *R* to obtain the posterior distribution for *R*. Under the true distribution, the reproduction number was well estimated, in general, as expected.Across the 24 datasets, the obtained 95% HPD interval captured the true value of *R* in 22 datasets (Figure 5A). The two remaining datasets overestimated the reproduction number. The medians of the posterior distribution was slightly higher, with a mean of 3.149 (Figure 5B), but was not significantly different from the true value (one-sample Wilcoxon rank test, *p* = 0.08).

Consistent with the results from the coalescent and birth-death models, the misspecified exponentially distributed generation interval parameterized with the true mean leads to an underestimation of the reproduction number. Only 11 datasets had their 95% HPD interval capturing the true value, and the underestimation of *R* was observed in 13 datasets (Figure 5A). The median was significantly lower than the true value (one-sample Wilcoxon rank test, *p <* 0.001), and the mean value was 2.580 across 24 datasets (Figure 5B). The size of the 95% HPD interval under the exponential distribution was significantly smaller (Mann-Whitney U rank test, *p <* 0.001; Figure 5C). This shows that the underestimation of *R* and the uncertainty in the estimates under exponential distribution were observed across tree models, suggesting the importance of the generation interval distribution in phylodynamic inferences.

### 3.2 Underestimation of *R* can be explained by the *R* − *r* relationship

The analyses in Section 3.1 indicated that *R* was systematically underestimated when the distribution of the generation interval was misspecified with an exponential distribution rather than with the true distribution that has a smaller variance. Similar underestimation is also observed in case-based inferences relying on the growth rate to estimate *R*, as the misspecified variation affects the *r* − *R* relationships (5; 18). The exponential distribution, which has higher variance than the true distribution, has a higher density at a shorter generation interval, which implies that infections tend to occur earlier than in the true distribution (orange and blue lines in Figure 1B, also see (5)). These infections drive the faster growth of the epidemic and, therefore, lead to a higher growth rate under a given reproduction number (5). Namely, under the misspecified exponential distribution, the estimated *r* corresponds to a lower *R* compared to the true distribution with smaller variance (orange and blue lines in Figure 1C, also see (5; 18; 19)).

In the following section, motivated by case-based inferences, we investigate whether the growth rate can explain the underestimation of *R* in phylodynamic inferences by performing phylodynamic analyses with a new exponential distribution (green line in Figure 1C) that is adjusted to match the true intrinsic growth rate. This new exponential distribution has a longer mean generation interval 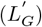 of 9.470 days, compared to the original mean generation interval of 6.963 days. If the growth rate is the main driver of the underestimation, estimates under the exponential distribution with the adjusted mean will perform better than those with the true mean.

#### 3.2.1 Coalescent exponential model

We re-calculated the reproduction number from the estimated growth rate (2) assuming an exponential distribution with the adjusted mean 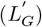. As expected, the underestimation observed with an exponential distribution parameterized with the same mean as the true distribution is no longer observed. Among the 24 analyzed datasets, the true *R* was recovered in 22 datasets, as under the true generation interval. The datasets that failed to recover the true *R* were the same datasets that failed to recover the true *r* and true *R* under the true generation interval distribution. The median of the posterior distribution of *R* was slightly higher than the true value, with a mean of 3.194 on average (one-sample Wilcoxon rank test, *p <* 0.001, Figure 3B).

The median of the estimated *R* was not significantly different from that under the true generation interval distribution (Mann-Whitney U rank test, *p >* 0.05). This further supports that the underestimation observed under exponential distribution with true mean is due to the *r* − *R* relationship (Figure 1C). Unlike the exponential distribution with a true mean that has a higher growth rate, the new exponential distribution with an adjusted mean has a comparable growth rate, as the adjusted mean is longer than the true mean. Therefore, the new distribution does not lead to a lower *R* estimate to compensate for the higher growth, as observed under the exponential distribution using the true mean.

The size of the 95% HPD interval was comparable to those under the true distribution (Mann-Whitney U rank test, *p >* 0.05). However, the mean of the interval size was slightly lower under the exponential distribution with adjusted mean, with the mean of 1.602, compared to the mean of 2.191 under the true distribution (Figure 3C). Again, this can be explained with the slope for *R* = *f* (*r*) under the true and exponential distribution (Figure **??**C). The *R* = *f* (*r*) function for the true distribution and the exponential distribution with the adjusted mean intersects at the true *R*. However, as *f* (*r*) is a concave function, for *r* that is less than the true value, the *R* for the true distribution is lower than that for the exponential distribution with adjusted mean. Likewise, for *r* that is greater than the true value, the converted *R* is higher under the true distribution. Together, these lower *R* for *r < r*_*true*_ and greater *R* for *r > r*_*true*_ leads to the wider 95% HPD interval for the true distribution.

#### 3.2.2 Birth-death model

Similarly, under the birth-death model, the exponential distribution with the longer mean resulted in higher *R* estimates that were closer to the true value. Across 24 datasets, the true *R* was recovered in 20 datasets (Figure 4A). Among the four datasets that failed to recover *R*, two underestimated the true value, and two overestimated the true value. The median of the posterior distribution of *R* was 2.991 on average, which was very close to the true value of 3.0 (one-sample Wilcoxon rank test, *p* = 0.843). The median estimates of *R* were not significantly different from those under the true distribution (Mann-Whitney U rank test, *p >* 0.05; Figure 4B). As in the exponential growth coalescent model, using the adjusted mean could recover the true reproduction number, suggesting the importance of the growth rate in the estimation of the reproduction number. Compared to the true distribution, the 95% HPD interval size under the exponential distribution with adjusted mean is not significantly different. There was no significant difference between the 95% HPD interval size under the true distribution and exponential distribution with adjusted mean.

Unlike the coalescent exponential growth model, the birth-death model estimates the reproduction number directly without converting from the growth rate. To better understand whether the generation interval distribution affected the estimation of the growth rate, we further investigated the latent variables, in particular, the number of infected individuals over time. We found that the number of infected individuals increased at comparable rates, resulting in similar growth rate estimates despite differences in other parameters (Figure 6 upper panels). The underestimation of the reproduction number while the growth rates remain consistent across models suggests that the generation interval distribution affects the estimation of *R* primarily through the *r* − *R* relationship, as in the coalescent exponential growth model.

**Figure 6.**
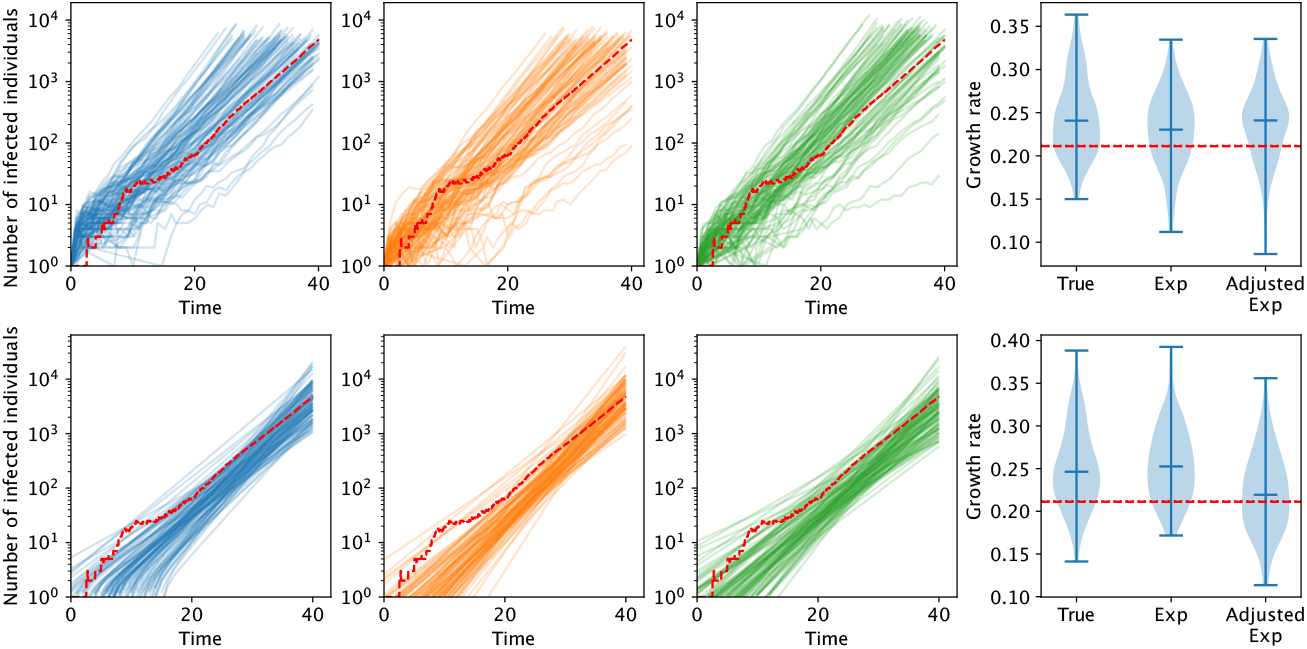
Example trajectory of the number of infected individuals and growth rate. The number of infected individuals over time is sampled during MCMC chains for the birth-death model (upper panels) and coalescent with the complex dynamics model (lower panels) for a simulation replicate. Each line represents a trajectory from one chain, and 100 chains are shown. Red dashed lines show the true simulated trajectory. For the true distribution (shown in blue), the number of infected individuals is obtained from the number of individuals in the E and I compartments. For the exponential distribution with true and adjusted mean (shown in orange and green), the number of infected individuals is obtained from the number of individuals from the I compartment. The rightmost panels show the distribution growth rate calculated from the sampled trajectory. The vertical line inside the violin plot indicates the median of the distribution, and the red vertical line indicates the true growth rate calculated from the true reproduction number.

#### 3.2.3 Coalescent model with complex dynamics

Consistent with other models, the estimated *R* values were higher and closer to the true value if the adjusted mean 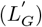 was used with the exponential distribution. The true *R* was recovered in 22 datasets, except for two datasets that overestimated *R* (Figure 5 A). The median of the *R* estimates was not significantly different from the true value (one-sample Wilcoxon rank test, *p* = 0.16) with a mean value of 3.08. The distribution of the median estimates from 24 datasets was not significantly different from those under true distribution (Mann-Whitney U rank test, *p >* 0.05). The size of the 95% HPD interval was significantly lower than those from the true distribution (Mann-Whitney U rank test, *p <* 0.05). Again, the underestimation of the *R* under the exponential distribution was not observed with a longer mean generation interval, which resulted in a comparable growth rate, suggesting that the underestimation is mainly governed by the *r* − *R* relationship.

As in the birth-death model, the latent variables for epidemiological dynamics were investigated. The number of infected individuals over time showed similar patterns across generation interval distribution, and the growth rate estimated from each trajectory also showed similar distributions (Figure 6 lower panels). This further emphasizes the role of the *r* − *R* relationship in the phylodynamic estimation of *R*.

## 4 Discussion

In the analysis of epidemiological case data, it is well known that the generation interval distribution shapes the relationship between the epidemic growth rate and the reproduction number (3; 5). However, the impact of this relationship on phylodynamic analyses has been less well-studied. In this study, we focus on the impact that a commonly used, but oftentimes misspecified, exponentially distributed generation interval distribution has on the estimation of *R*. We demonstrate that assuming an exponentially distributed generation interval when the true generation interval distribution has a smaller variance leads to a systematic underestimation of the reproduction number and erroneously high confidence in this underestimate.

Our results were consistent with those from case-based inferences. In case-based inferences, the growth rate can be estimated first from the incidence data and then used to calculate the reproduction number from the estimated growth rate (3). In the exponential growth coalescent model, just as in case-based inferences, the growth rate is estimated first, and then the reproduction number is estimated. Therefore, a similar bias is expected under the exponential growth coalescent model. However, even when the generation interval is incorporated through other model components in the BDMM and PhyDyn model, we observed the underestimation of *R* under the exponential distribution.

In our findings, the exponential distribution with adjusted mean successfully recovered the true growth rate. Although this might suggest that the underestimation could be explained by matching the growth rate, we emphasize that this approach is not applicable to real-world data analysis. In our analyses, we could calculate the adjusted mean generation interval based on the true reproduction number. However, in the real world, we do not know the true reproduction number; it is the parameter we aim to estimate. Therefore, our results demonstrate the role of growth rate in the underestimation of the reproduction number rather than suggest a way to correct the bias in existing inference approaches.

Despite its importance in inference, the generation interval is a rarely observed quantity, as it is very hard to know exactly when one is infected. Therefore, the exact distribution for the generation interval is also rarely known. However, (5) suggested that approximating the generation interval distribution as a gamma distribution could be parameterized based on the mean and standard deviation or through maximum-likelihood estimation. This provides a way to incorporate the generation interval into inference approaches and use the known *r* − *R* relationship for gamma distribution (3). In phylodynamic analyses, however, we are not aware of any package that allows for gamma-distributed generation intervals in inference. Although the exposed compartment can generate the gamma distribution when the duration in exposed and infectious compartments are similar, this represents very limited cases. Moreover, having multiple compartments can significantly slow down phylodynamic analyses, limiting its usability. Therefore, developing a flexible phylodynamic inference approaches or extensions that can incorporate more flexible generation interval distributions would be valuable for future research.

