## Supplementary Materials for "Common misspecification of the generation interval leads to reproduction number underestimation in phylodynamic inference"

### Supplementary information

**Table S1.** Parameters and priors for BDMM-Prime analyses

| **Parameter** | **Prior** | **Unit** |
|---|---|---|
| clockRate | Unif(0, Infinity) | |
| originBDMMPrime ($T_0$) | Unif(0, Infinity) | days |
| **Assumption 1: True distn. from EI model** | | |
| birthRateAmongDemesCanonical.E_to_I ($\beta$) | Unif (0.3368, 3.3684) | days$^{-1}$ |
| birthRateAmongDemesCanonical.I_to_E | Fixed at 0 | days$^{-1}$ |
| birthRateCanonical.E | Fixed at 0 | days$^{-1}$ |
| birthRateCanonical.I | Fixed at 0 | days$^{-1}$ |
| deathRateSPCanonical.E | Fixed at 0 | days$^{-1}$ |
| deathRateSPCanonical.I ($\delta$) | Fixed at 1/3 | days$^{-1}$ |
| samplingRateSPCanonical ($\psi$) | Fixed at 0.0015 | days$^{-1}$ |
| removalProbCanonical | Fixed at 1 | NA |
| migrationRateSPCanonical.E_to_I ($\gamma$) | Fixed at 1/4 | days$^{-1}$ |
| migrationRateSPCanonical.I_to_E | Fixed at 0 | days$^{-1}$ |
| startTypePriorProbs.E | Fixed at 0 | NA |
| startTypePriorProbs.I | Fixed at 1 | NA |
| **Assumption 2: Exp. distn. with original mean** | | |
| birthRateCanonical ($\beta$) | Unif(0.1436, 1.4362) | days$^{-1}$ |
| deathRateSPCanonical ($\delta$) | Fixed at 0.1421 | days$^{-1}$ |
| samplingRateSPCanonical ($\psi$) | Fixed at $1.496 \times 10^{-3}$ | days$^{-1}$ |
| removalProbCanonical | Fixed at 1 | NA |
| **Assumption 3: Exp. distn. with adjusted mean** | | |
| birthRateCanonical ($\beta$) | Unif(1.056, 1.0559) | days$^{-1}$ |
| deathRateSPCanonical ($\delta$) | Fixed at 0.1045 | days$^{-1}$ |
| samplingRateSPCanonical ($\psi$) | Fixed at $1.100 \times 10^{-3}$ | days$^{-1}$ |
| removalProbCanonical | Fixed at 1 | NA |

**Table S2.** Parameters and priors for PhyDyn analyses

| **Parameter** | **Prior** | **Unit** |
|---|---|---|
| clockRate | Unif(0, Infinity) | |
| **Assumption 1: True distn. from EI model** | | |
| $\beta$ | Unif (0.3368, 3.3684) | days$^{-1}$ |
| $\gamma$ | Fixed at 1/4 | days$^{-1}$ |
| $\delta$ | Fixed at 1/3 | days$^{-1}$ |
| $\psi$ | Fixed at 0.0015 | days$^{-1}$ |
| $E_0$ | Fixed at 0 | |
| $I_0$ | Fixed at 1 | |
| **Assumption 2: Exp. distn. with original mean** | | |
| $\beta$ | Unif(0.1436, 1.4362) | days$^{-1}$ |
| $\delta$ | Fixed at 0.1436 | days$^{-1}$ |
| $I_0$ | Fixed at 1 | |
| **Assumption 3: Exp. distn. with adjusted mean** | | |
| $\beta$ | Unif(0.1056, 1.0559) | days$^{-1}$ |
| $\delta$ | Fixed at 0.1056 | days$^{-1}$ |
| $I_0$ | Fixed at 1 | |
